# OneGrow: Unified Temporal Plant Image and Mask Generation

**DOI:** 10.64898/2026.09.11.750899

**Authors:** Mike Boss, Michele Volpi, Lukas Roth

## Abstract

Image-based crop phenotyping benefits from image series that capture plant development together with organ-level labels. Such paired data are limited because organ annotation is expensive, and following the same plants over time requires repeated, registered imaging. Existing generative models for plants either synthesize temporal imagery without structural labels or generate labeled images without a temporal dimension. We introduce *OneGrow*, a latent flow-matching model that jointly models wheat images and their organ-segmentation masks over time. Images and masks share a single frozen image autoencoder. A reveal specifies which content is observed context, so the same model covers tasks such as mask-to-image synthesis, image-to-mask segmentation, and temporal forecasting. For sequences longer than the training window, a sliding-window roll-out generates each new image from the preceding frames, keeping long sequences temporally consistent. We train jointly on a large single-frame wheat dataset and a multi-year temporal dataset, using pseudo-labels from a pretrained segmentation model. We evaluate segmentation and image quality against held-out references and assess the multi-task and temporal behavior qualitatively.

## 1 Introduction

Image-based phenotyping derives plant traits from field imagery and is widely used to study crop development [9, 33, 41, 48]. Such imagery becomes considerably more useful when it carries two additional kinds of information, both of which are costly to obtain. The first is organ-level structure, provided by segmentation masks. Their annotation is labor-intensive, so labeled data cover only a small fraction of collected imagery [48]. The second is temporal development, captured by imaging the same plots repeatedly across a season, which few platforms undertake [41]. Imagery that provides both at once remains scarce.

Generative models may expand such data. Diffusion and flow-matching models synthesize images across many domains [16, 20, 28, 40], and several have been adapted to plants. Existing plant models either predict imagery at unobserved time points without producing structural labels [14, 15, 47] or generate labeled image-mask pairs for a single frame [18]. To our knowledge, no plant-focused model jointly generates organ-labeled imagery over time.

We introduce *OneGrow*, a latent flow-matching model of wheat images and their organ masks over time. It builds on unified generative models that give each input and output its own flow time, a per-view position on the path from noise to clean data, so either can serve as context or a target [26]. Related methods jointly generate images and segmentations for single frames [38] or video with dense labels [51, 53]. OneGrow targets sparse seasonal observations of wheat canopies and jointly trains on a single-frame source with broad appearance variation and a temporal source that follows plots across years. It therefore places weaker demands on data availability than models that require densely sampled, fully labeled sequences.

OneGrow encodes RGB images and color-coded masks with the same frozen image autoencoder [26, 38]. It represents each window as a collection of views indexed by modality and frame. A *reveal* sets which views, and within a view which patches, are given as observed context and which content the model generates. The resulting objective covers image-mask tasks such as segmentation, synthesis, and forecasting. Training masks are pseudo-labels from a pretrained segmentation model, which makes both wheat sources available for joint training. A sliding-window roll-out generates frames beyond the frame count used in training, so image sequences stay temporally consistent over longer horizons.

Our contributions are:

- OneGrow, a unified latent generative flow-matching formulation for wheat images and organ masks over time.
- Joint training on a diverse single-frame wheat source and a multi-year temporal source using pseudo-labels from a pretrained segmentation model.
- A reveal mechanism over a (modality, frame) view collection that specifies tasks such as segmentation, synthesis, forecasting, and spatial completion. The reveal is recorded per patch, so observed context can be spatially partial.

## 2 Related Work

### Diffusion and flow-matching generative models

Diffusion and flow-matching models synthesize images by learning to transform noise into data. Diffusion models reverse a gradual noising process [20], while flow matching learns a velocity field that transports noise to data along a prescribed path [28, 29]. Both families commonly operate in the compressed latent space of a pretrained autoencoder, which lowers the cost of high-resolution synthesis [25, 40]. Recent large text-to-image models adopt the flow-matching objective [16]. OneGrow applies latent flow matching with a transformer backbone to joint image-mask temporal modeling.

### Unified and multi-task generation

Unified generative models avoid training a separate network for every conditioning signal. OneDiffusion [26] represents inputs and outputs as views carried at different flow times, so each view can act as context or a target. UniGS [38] applies this formulation to an image and its segmentation, encoding a color-coded mask with the same image autoencoder. Panoptic Diffusion Models also co-generate an image and a panoptic segmentation map [30]. Related video models jointly generate dense labels, depth, and frames, or select each modality as context or a target [6, 51, 53]. These methods target general scenes and densely sampled video. OneGrow instead models sparse seasonal wheat observations and encodes the organ segmentation masks as color-coded RGB images through a 2D image autoencoder.

### Masked and per-token generative modeling

Masked prediction provides a useful training signal for recognition and generation [4, 17]. Diffusion Forcing [5] assigns independent noise levels to tokens, which unifies next-token prediction and full-sequence diffusion. OneGrow sits between these and the per-view scheme above. Its reveal is recorded per patch, so observed context can be spatially partial rather than whole-view [26]. Its generated patches all share one flow time rather than independent ones.

### Plant imaging and phenotyping

Image-based phenotyping spans many crops and imaging tasks, and deep learning now drives much of this analysis [33]. Semantic segmentation of field imagery is a central tool [27], with dedicated benchmarks in the agricultural domain [49]. For wheat, benchmarks range from head detection [9] to organ-level semantic segmentation [48], while field-phenotyping platforms add image sequences of plant canopy instances over multiple years [41]. OneGrow uses the organ-segmentation data, pseudo-labeled by the challenge-winning segmentation model [3], together with a temporal platform that supplies aligned windows for generative modeling.

### Generative models for plants and crops

Prior plant generation follows two main directions. Conditional generative adversarial networks (GANs) predict images at unobserved times from earlier frames [13–15, 32]. Recent diffusion models extend temporal generation to crop video [47]. Other generative models condition plant growth on environmental drivers [10]. Surveys cover this growing literature [11, 22]. A second direction generates labeled image-mask pairs for segmentation training in agriculture [18] and other domains [34, 38, 45, 50, 55]. These approaches typically address a single frame. OneGrow combines temporal image generation with joint image-mask modeling for wheat.

### Crop-growth models

Process-based crop-growth models simulate development through traits and state variables rather than imagery. APSIM [21] and DSSAT [24] are representative examples. Because they produce traits rather than spatial structure, their output cannot directly condition OneGrow. Functional-structural plant models and procedural generators instead build explicit plant geometry, from which organ masks can be rendered [1, 2, 46]. OneGrow can translate such mask sequences into images conditional on temporal context.

## 3 Method

### 3.1 Overview and problem setting

OneGrow models wheat images together with their organ masks over time. Its training data pairs every image with a mask and groups frames of the same plot into a short growth window (Fig. 1(A)). The single-frame source forms windows of length one, while the temporal source supplies multi-frame windows (Section 4.1). Each frame carries one or more *modalities*. We use two, an RGB image and an organ-segmentation mask. The window is also accompanied by conditioning metadata, namely the acquisition domain and per-frame photometric statistics for exposure and color (Section 3.6).

**Figure 1:**
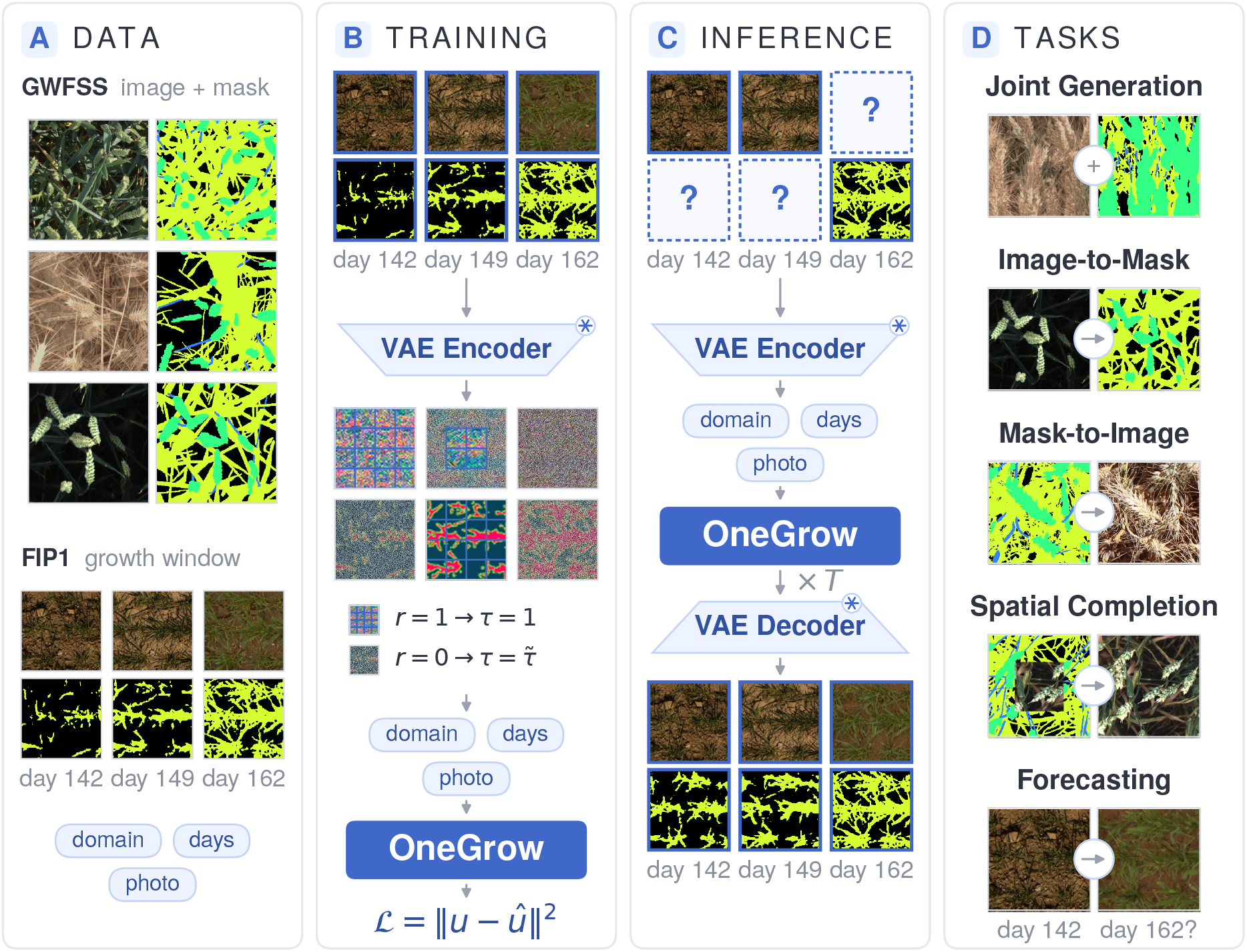
Overview of OneGrow. **(A)** Training data comes from two paired image-and-mask sources: single-frame GWFSS samples spanning several domains and multi-frame FIP1 growth windows, together with the acquisition and appearance metadata used for conditioning. **(B)** At training time, the image and mask views of a window are encoded by one shared frozen autoencoder, the reveal marks each patch as clean context or as generated, and one transformer regresses the joint velocity field under the flow-matching loss. **(C)** At inference, the observed views are revealed as clean context and the remaining views are generated by integrating the flow over several steps, then decoded to pixels. **(D)** One single trained model covers many tasks, including joint image-mask generation, image-to-mask segmentation, mask-to-image synthesis, spatial completion, and temporal growth forecasting. Each task is selected by a different reveal configuration.

OneGrow generates a window in one shared latent space (Fig. 1(B)). A single frozen autoencoder encodes every image and mask to latent patches, so the model needs no modality-specific encoder. One flow-matching transformer processes all patches of the window at once and predicts their velocity jointly. A reveal marks which content is given as observed context and which the model generates. Which views are observed selects the task, and the reveal is recorded per patch so a single view can hold observed and generated patches at once. The reveal is the only part of the input that changes between tasks.

At inference, the reveal fixes the observed views, and the transformer turns noise into the remaining views, which the autoencoder decodes to pixels (Fig. 1(C)). The choice of reveal selects the task, so one trained model performs many reveal-defined tasks (Fig. 1(D)).

### 3.2 Shared image-mask latent representation

Images and masks use one shared representation, following unified image-and-label models [26, 38]. Both modalities are encoded by the same frozen, pretrained image variational autoencoder (VAE) [25, 40]. The encoderℰ maps an *H× W* image *x* to a latent *z* = ℰ (*x*) *∈* ℝ^*C×h×w*^, and the decoder *D* maps a latent back to image space, where *C* denotes the number of latent channels and *h* and *w* denote the spatially downsampled latent height and width, respectively.

A segmentation mask is a per-pixel class map *y∈{* 0, …, *K*− 1*}*^*H×W*^ over *K* semantic classes. For wheat, these are background, head, stem, and leaf. To encode it with the same image autoencoder, we *color-code* the mask into an RGB image using a fixed palette *π* : *{*0, …, *K*−1*}→* [0, 1]^3^, giving *x*^mask^ = *π*(*y*) [38]. The palette colors stay separable through the autoencoder round-trip, so after decoding each pixel is assigned to its nearest palette color. Encoding and decoding a color-coded mask recovers it at a mean intersection-over-union of 0.990, so the representation is not what limits the segmentation results of Section 5.1. Image and mask latents differ only in their *modality* index.

### 3.3 The window as a view collection

The window is indexed by *views*. Let *V ⊆ {*0, …, *M*− 1*}× {*0, …, *F*− 1*}* denote the present views, where *m*=0 denotes RGB images and *m*=1 denotes masks. A full paired window has *M* =2 modalities and *V* =|*V*| = *MF* views. A view *v*=(*m, f*) holds a latent *z*_*v*_ ℝ^*C×h×w*^, split into *P* non-overlapping *p× p* patches.

Each view has explicit modality and frame coordinates. For temporal windows, frame coordinates are ordered by elapsed acquisition time *t*_*f*_ . Frames without a known acquisition time use a learned null-time embedding. Representing the input as a collection indexed by these coordinates lets the model take a variable number of views, including partial and non-contiguous subsets, without a fixed image-mask layout.

### 3.4 Flow-matching architecture

OneGrow is a flow-matching diffusion transformer (DiT) [28, 29, 36]. It connects a clean latent *z*, the encoding of an observed image or mask, and Gaussian noise *ϵ ∼ N* (0, *I*) with the linear interpolant

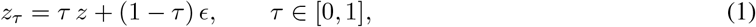

where *τ* is the *flow time, τ* =1 is clean data, and *τ* =0 is noise. The corresponding velocity target is

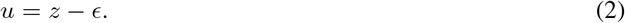

The reveal is set per patch, so clean context and generated content can coexist in one view. Within a window the flow time takes only two values, *τ* =1 on revealed patches and a single shared value on the generated ones.

The network *u*_*θ*_ predicts all patch velocities jointly. Each latent patch becomes a token augmented with its modality and frame coordinates. The transformer uses rotary position embedding (RoPE) over modality, time, and spatial coordinates, while flow time enters through adaptive layer-normalization modulation [36, 44]. Each block applies query-key normalization for training stability [12] and uses SwiGLU feed-forward layers [42]. A shared linear head maps tokens back to velocity patches.

### 3.5 Reveal-based multi-task objective

The observed portion of a window is encoded by a binary reveal *r∈ {* 0, 1*}* ^*V × P*^ over views and patches. Here *r*_*v,j*_=1 marks patch *j* of present view *v∈ V* as clean context at *τ* =1, while *r*_*v,j*_=0 marks a generated patch. Its flow time is

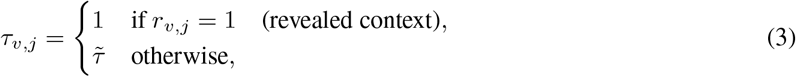

where 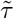 is drawn once per window from a logit-normal schedule [16]. Thus, the reveal determines which content is given, while the shared scalar 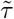 determines the flow state of all generated patches. This separates spatially partial context from the noise schedule.

The model regresses the velocity target of generated patches using the reveal-weighted mean squared error (MSE)

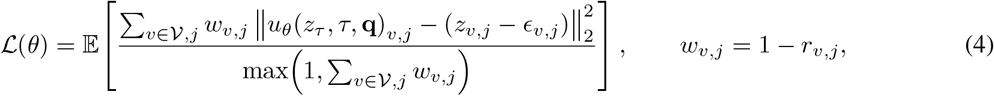

where **q** denotes the conditioning tokens and the expectation is over the data, the noise *ϵ*, the reveal *r*, and 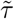. The weight *w*_*v,j*_ is zero on revealed patches, so only generated patches contribute to the loss.

A reveal configuration specifies a task, and most tasks are selected at the level of whole views. Revealed image views and generated mask views define image-to-mask segmentation (Fig. 2). Reversing these roles defines mask-to-image synthesis (Fig. 3). Revealing earlier frames defines forecasting (Fig. 4). An unrevealed window supports joint image-mask generation, and the temporal roll-out reveals prior image views together with the target mask to generate the target image. Spatial completion is the exception, where a spatially partial reveal splits one view into observed and generated patches (Fig. 5).

**Figure 2:**
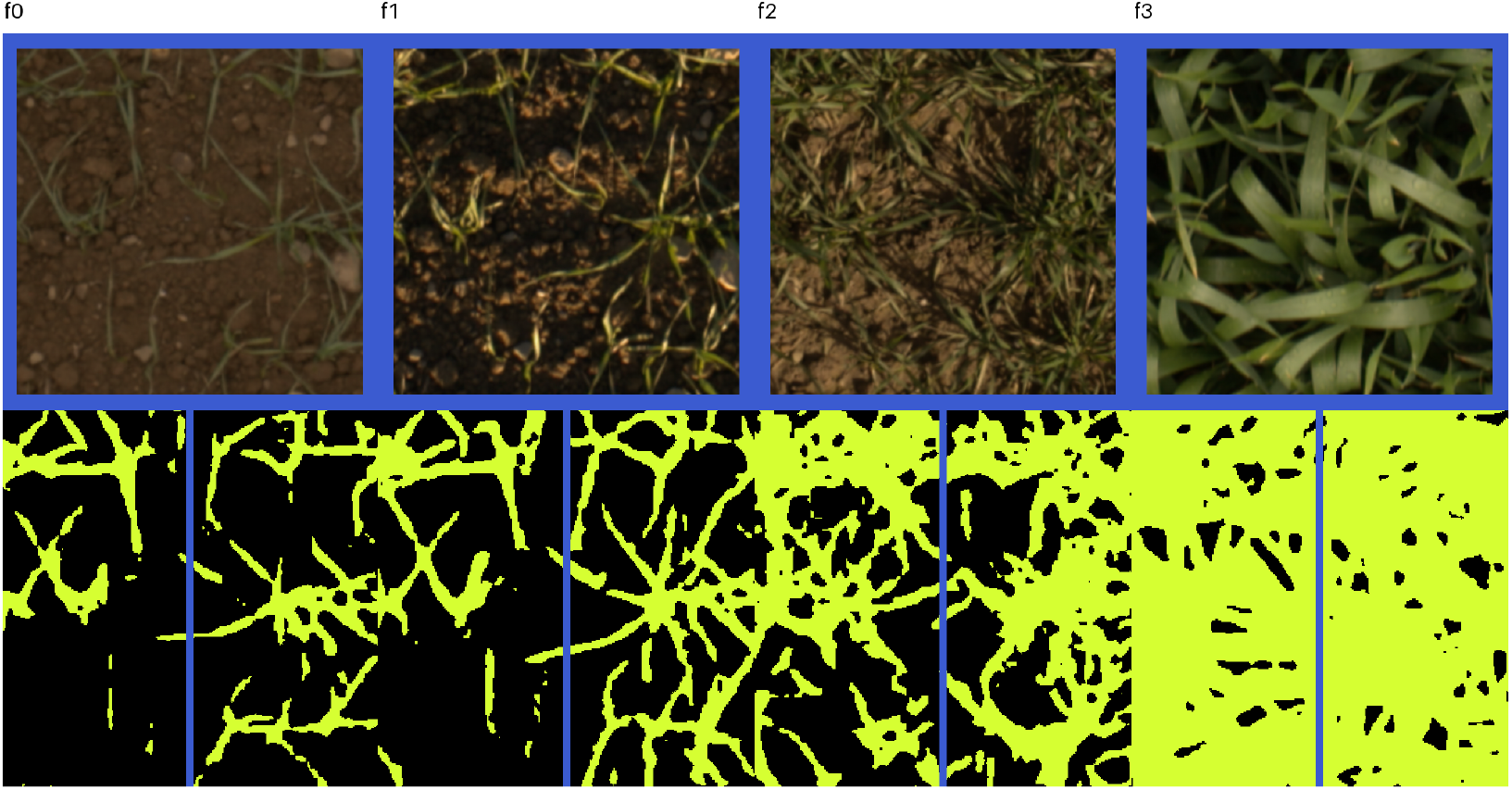
Image-to-mask segmentation on a FIP1 growth window. The image views (*top row*) are revealed as clean context, and OneGrow generates the organ masks (*bottom row*). This early-season FIP1 window contains only two classes, background (black) and leaf (yellow-green).

**Figure 3:**
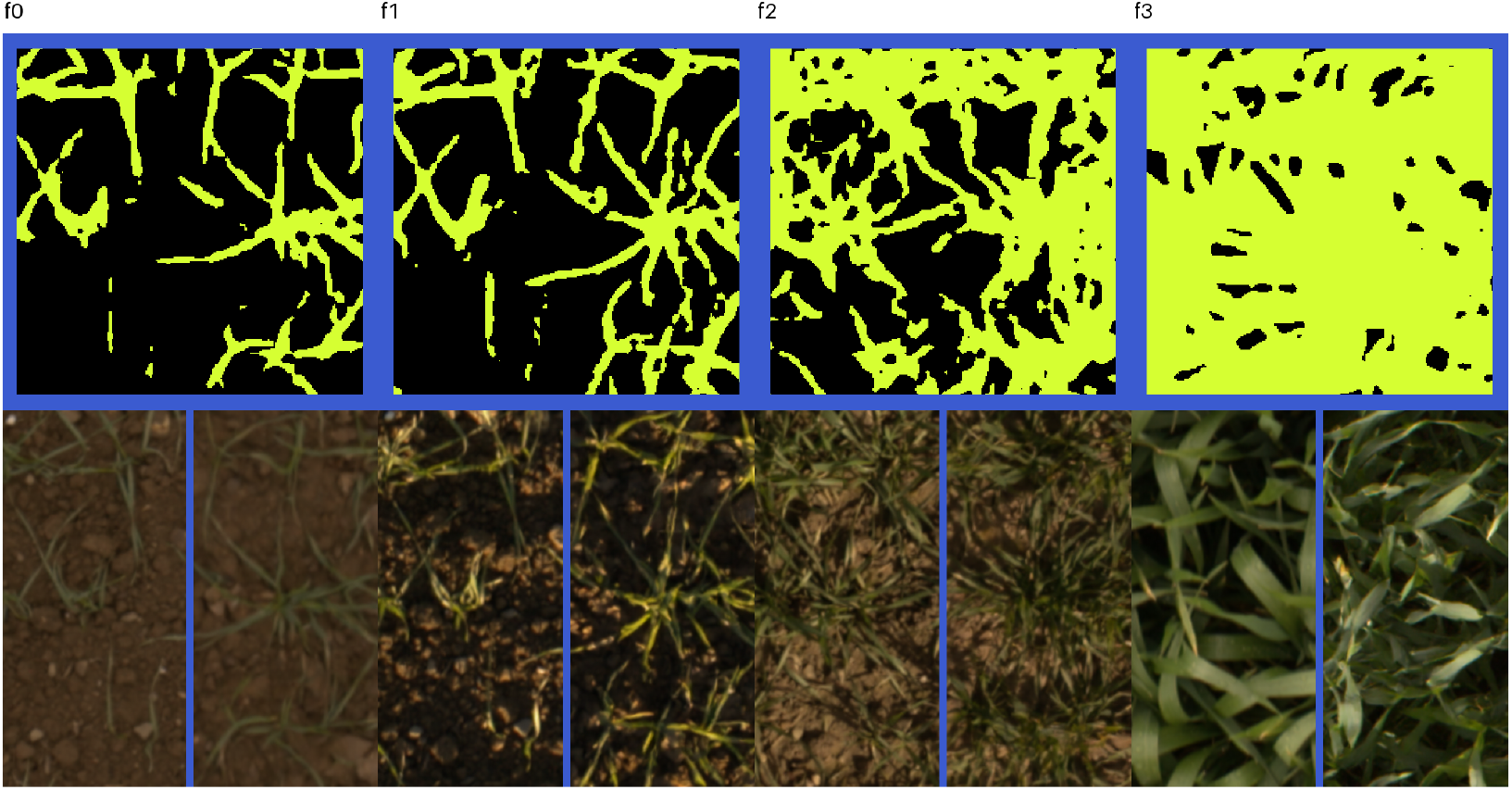
Mask-to-image synthesis on a FIP1 growth window. The organ masks (*top row*) are revealed as clean context, and OneGrow generates the images (*bottom row*). The seam falls mid-cell, so continuity between the two halves is visible.

**Figure 4:**
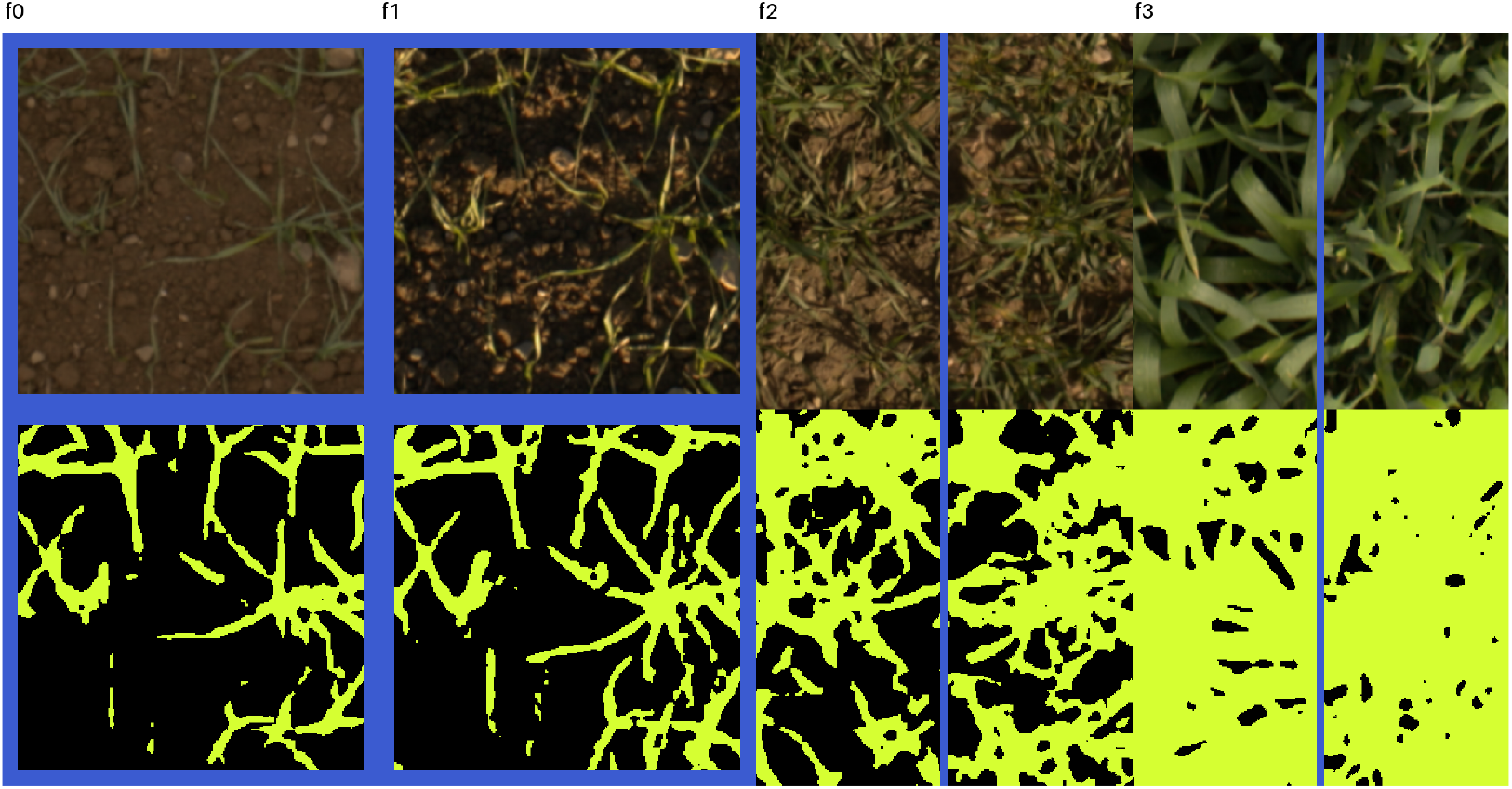
Temporal forecasting on a FIP1 growth window. The image (*top row*) and mask (*bottom row*) of the first two frames are revealed as clean context, and OneGrow forecasts the later frames.

**Figure 5:**
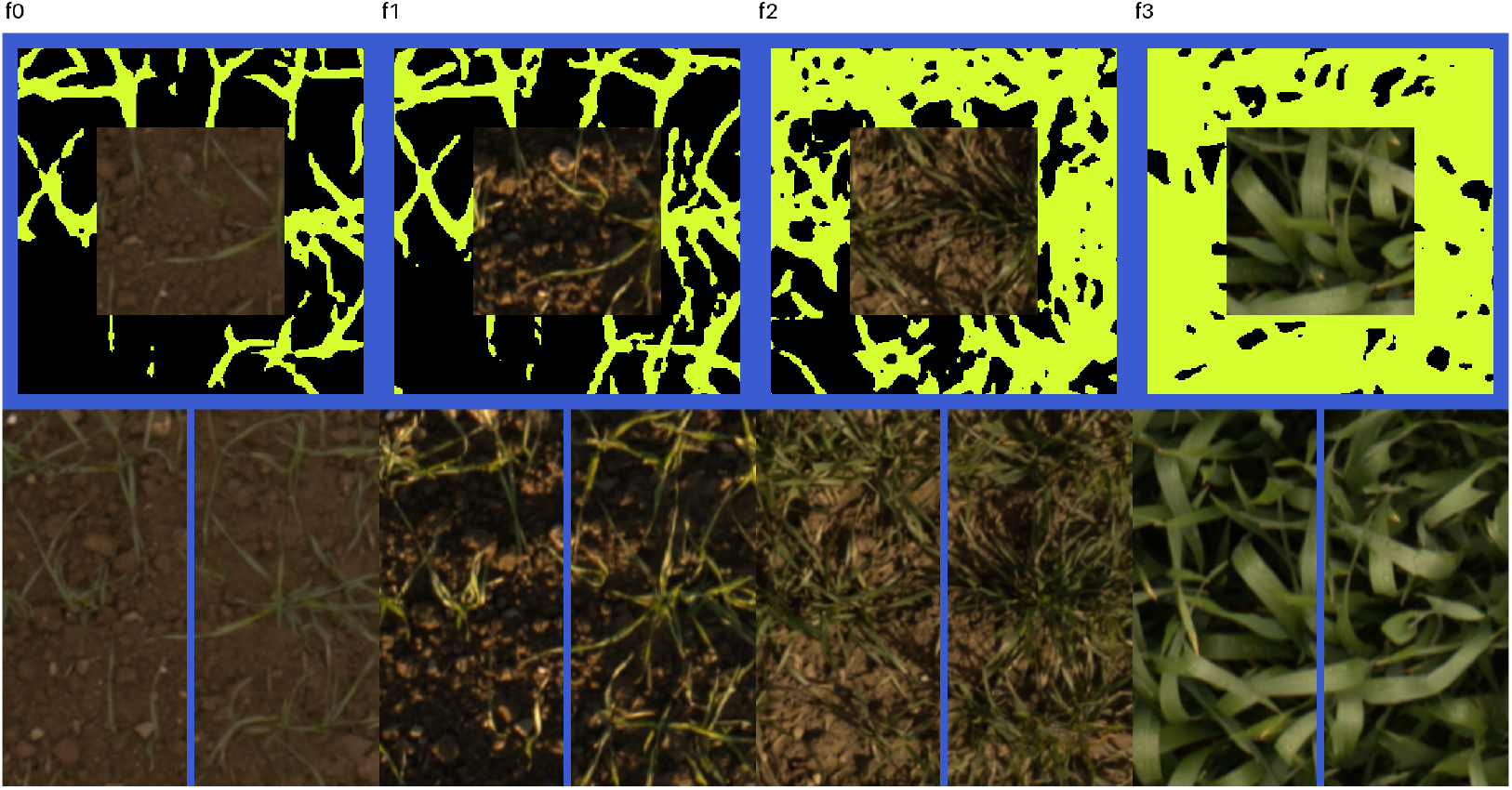
Mask-guided spatial completion on a FIP1 growth window. The full organ mask with a central image patch composited into its center (*top row*) is revealed as clean context, and OneGrow outpaints the surrounding image (*bottom row*).

Figures 2 to 5 show these configurations on FIP1 growth windows. In each, columns *f*_0_ to *f*_3_ are frames of one plot ordered by acquisition time, content outlined in blue is revealed as clean context, and every generated cell is split down the middle into the held-out reference (left half) and the generated content (right half). A generated half is a sample and need not match the reference exactly.

During training, reveal configurations are drawn from a fixed non-uniform distribution over task families that emphasizes the target tasks of mask-conditioned synthesis and forecasting while still covering the full range of image-mask relationships.

Training also applies per-batch *view dropout*, which drops a shared random subset of views. The model therefore sees variable, sometimes unpaired image and mask context during training. Validation and evaluation use the complete available view set.

### 3.6 Conditioning

Beyond reveal context, the model accepts optional per-window metadata. Each conditioner is represented by a *register token* with a learned null embedding. The reported model conditions on the acquisition *domain* and on per-frame *photometric* statistics for exposure and color. Conditioners are independently dropped during training for classifier-free guidance (CFG) [19].

### 3.7 Sampling and temporal roll-out

At inference, generated patches start from noise and revealed patches remain pinned to their clean latent. The flow is integrated toward *τ* =1 with CFG over the available conditioners [16, 28]. The model can sample any reveal configuration supported by the supplied views.

#### Temporal roll-out beyond the training window

A sequence can be longer than the window seen in training. We extend the model with a stride-one sliding window, rolling it out autoregressively beyond its training horizon [5]. Each step reveals the already-generated frames of the overlap as context and generates the next frame, so every frame is conditioned on its predecessors and the sequence stays temporally consistent. The reveal can additionally fix any further available views, such as a per-frame mask that guides the generated image or observed early frames that seed the roll-out. Without such views, the same procedure rolls out unconditionally. The roll-out reuses the trained reveal mechanism and adds no training objective.

## 4 Experimental Setup

### 4.1 Datasets and pseudo-labeled masks

Training uses two complementary wheat-imaging sources. The *GWFSS* dataset was released for a full semantic organ segmentation challenge [48]. Its labeled portion is small. The challenge released 99 images with human organ masks for training and about 64k unlabeled images for pretraining. The 1,096-image challenge test set was made public with human masks after the challenge closed. We pseudo-label these 64k images and train OneGrow on the resulting pairs. They span nine institutions and give broad appearance variation without temporal structure. We hold out the 1,096 labeled images for segmentation evaluation and never train OneGrow on them. The *FIP1* field-phenotyping dataset [41] supplies the temporal training signal, contributing high-resolution time series from approximately 4,000 winter-wheat plots acquired over six years. We form each plot sequence into windows of *F* =4 frames. The native frames are 960*×* 640. OneGrow takes a plot-aligned 256 *×*256 inner-plot crop of each frame and upsamples it to 512*×*512 before encoding.

The pseudo-labels come from the winning entry of the GWFSS segmentation challenge, a ViT-Adapter and Mask2Former network with a masked-image-modeling backbone [3, 7, 8, 37]. Its predictions are *pseudo-labels*, not human annotations [52]. We use them only for OneGrow training and retain the held-out human-annotated GWFSS masks for evaluation. These held-out masks are the challenge test set, which this segmentation model was ranked on but never trained on, so the segmentation evaluation stays independent of the pseudo-label teacher. For FIP1, the model runs tile-wise and blends per-class scores. Each image-mask pair is then encoded to latents as described in Section 3.2. FIP1 shares an acquisition domain with one GWFSS institution, which links the two sources through the domain conditioner.

### 4.2 Training details

We train one model jointly on both sources. GWFSS contributes single-frame windows and FIP1 contributes four-frame windows, both with pseudo-labeled masks. The two sources are sampled with equal probability. Both modalities use the frozen SD3.5 VAE [16] with 16 latent channels and eightfold downsampling, so a 512*×* 512 image becomes a 64*×*64 latent. The transformer has width 512, 16 blocks, 8 attention heads, and latent patch size 2, giving *P* =1024 patches per view. We optimize Eq. (4) with AdamW [31], using *β*_1_=0.9, *β*_2_=0.95, weight decay 0.01, a peak learning rate of 3*×* 10^−4^, linear warm-up, cosine decay, gradient clipping, and an exponential moving average for evaluation. Unless noted, sampling uses a midpoint ordinary differential equation (ODE) solver with 50 steps and guidance scale 2.

### 4.3 Tasks and metrics

We evaluate segmentation quantitatively and assess the remaining tasks qualitatively. *Segmentation* is measured by mean intersection-over-union between generated masks and held-out human-annotated GWFSS masks (Section 4.1). This evaluation measures organ recovery rather than agreement with pseudo-labels. FIP1 has no human annotations, so on FIP1 we report the same metric against the teacher pseudo-labels as a proxy. This measures agreement with the teacher rather than organ recovery, and is not directly comparable to the GWFSS numbers. We also measure *mask adherence*, the degree to which a mask-to-image generation respects the organ layout it was conditioned on, a property the realism metrics below do not assess. We quantify it through cycle consistency by re-segmenting the generated image and scoring the result against the conditioning mask (Section 5.1). *Image generation* is assessed against held-out reference frames with metrics suited to small reference sets and to imagery outside the ImageNet domain. We report the Fréchet distance in DINOv2 feature space (FD-DINOv2) [35, 43], the CLIP maximum mean discrepancy (CMMD) [23, 39], and paired Learned Perceptual Image Patch Similarity (LPIPS) [54].

## 5 Results

### 5.1 Segmentation

We first compare generated masks against held-out human-annotated GWFSS masks. OneGrow reveals image views and generates mask views. We report per-class and mean intersection-over-union and compare against the winning entry of the GWFSS segmentation challenge [3], applied to real images. We additionally report mask adherence (Table 1, last row).

**Table 1:** Image-to-mask segmentation and mask adherence, per-organ and mean intersection-over-union. Higher is better. The first two GWFSS rows predict a mask from a real image, for the challenge winner [3] and for OneGrow in its image-to-mask configuration. The third measures mask adherence by cycle consistency (Section 4.3). On GWFSS the reference is the held-out human-annotated masks (Section 4.1). FIP1 has no human annotations, so its row scores agreement with the teacher pseudo-labels and is not directly comparable to the GWFSS numbers.

| Dataset | Method | Head | Stem | Leaf | Background | mIoU ↑ |
| --- | --- | --- | --- | --- | --- | --- |
| GWFSS | Challenge winner | 0.847 | 0.488 | 0.833 | 0.791 | 0.740 |
|  | OneGrow (image-to-mask) | 0.811 | 0.412 | 0.805 | 0.758 | 0.697 |
|  | OneGrow (mask-to-image-to-mask) | 0.838 | 0.504 | 0.845 | 0.807 | 0.749 |
| FIP1 | OneGrow (image-to-mask) | 0.701 | 0.252 | 0.803 | 0.843 | 0.650 |

In its image-to-mask configuration, OneGrow reaches a mean intersection-over-union of 0.697, against 0.740 for the challenge winner, with the largest gap on the thin stem class (Table 1). Mask adherence reaches a mean intersection-over-union of 0.749, on par with the 0.740 the challenge winner achieves on real images. That 0.740 is a reference point for this scorer on real imagery rather than an upper bound on adherence. The cycle scores a generated image against the mask it was generated from, so the annotation ambiguity that limits segmentation of real imagery is absent. The challenge winner also produced the training pseudo-labels and performs the re-segmentation, so the value may favor appearances it segments reliably. Adherence stays below the challenge winner on the head class, at 0.838 against 0.847. On FIP1, where the references are teacher pseudo-labels, OneGrow reaches a mean intersection-over-union of 0.650 (Table 1). The challenge winner also produced the GWFSS pseudo-labels that OneGrow trains on, so its 0.740 indicates how accurate that training signal is. Its weakest class is stem, at 0.488, which is also the class where OneGrow is weakest. FIP1 has no human masks, so the accuracy of its pseudo-labels is unknown.

### 5.2 Image-generation quality

For mask-to-image synthesis, OneGrow reveals mask views and generates image views. We assess generated images against held-out imagery with the image-generation metrics of Section 4.3 and compare against the Mask-NN retrieval baseline (Table 2). On GWFSS, OneGrow attains a lower paired LPIPS than Mask-NN, while Mask-NN attains lower FD-DINOv2 and CMMD. Mask-NN returns real training images, so it scores well on FD-DINOv2 and CMMD, which compare sets of images rather than pairs. It cannot return the particular held-out frame it is scored against, which is what paired LPIPS measures. On FIP1, OneGrow attains a lower CMMD and a lower paired LPIPS than Mask-NN, and the two reach a comparable FD-DINOv2. The FIP1 scores use 250 held-out windows against 1,096 held-out images on GWFSS, so the set-level metrics carry more sampling noise there.

**Table 2:**
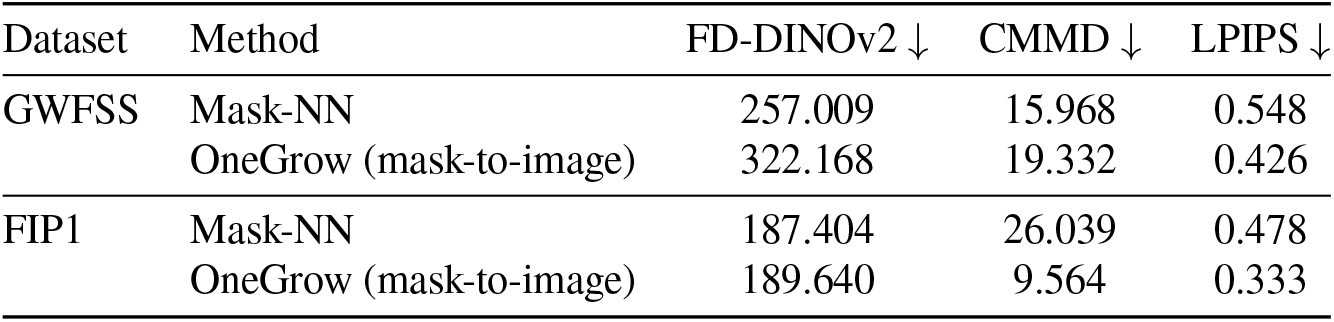
Mask-to-image generation quality on GWFSS and FIP1, against held-out reference imagery. Lower is better. *Mask-NN* is a training-free retrieval baseline that returns, for each test mask, the real training image whose mask latent is nearest. It conditions on the mask only, retrieves from the training split, and is scored against the held-out set.

### 5.3 Multi-task generation

We illustrate several image-mask relationships qualitatively on FIP1 growth windows. Tasks such as image-to-mask segmentation, mask-to-image synthesis, temporal forecasting, and mask-guided spatial completion use the same model and differ only in their reveal configuration. For image-to-mask segmentation, the generated organ masks follow the leaf and soil layout of the input and track the increasing canopy cover across the window (Fig. 2). For mask-to-image synthesis, the generated frames place vegetation and soil consistent with the conditioning mask and add plausible texture and illumination (Fig. 3). For temporal forecasting, the generated frames continue the growth trajectory toward denser canopy while remaining consistent with the revealed context (Fig. 4). For mask-guided spatial completion, the completed region matches the revealed image patch at its boundary and follows the surrounding mask (Fig. 5). Joint completion without a mask and the same tasks on single-frame GWFSS images are shown in Appendix A.

The same spatial-completion reveal lets the model grow images beyond its native window size. Starting from a single real image-mask tile, we outpaint half-overlapping windows outward and assemble a larger canvas of both modalities (Fig. 6). Each new window reveals its overlap with the already-generated canvas and jointly generates the remaining image and mask, so every seam falls on revealed content. The model thus produces large, spatially coherent image-mask pairs from a single seed tile, which can serve as additional labeled training data.

**Figure 6:**
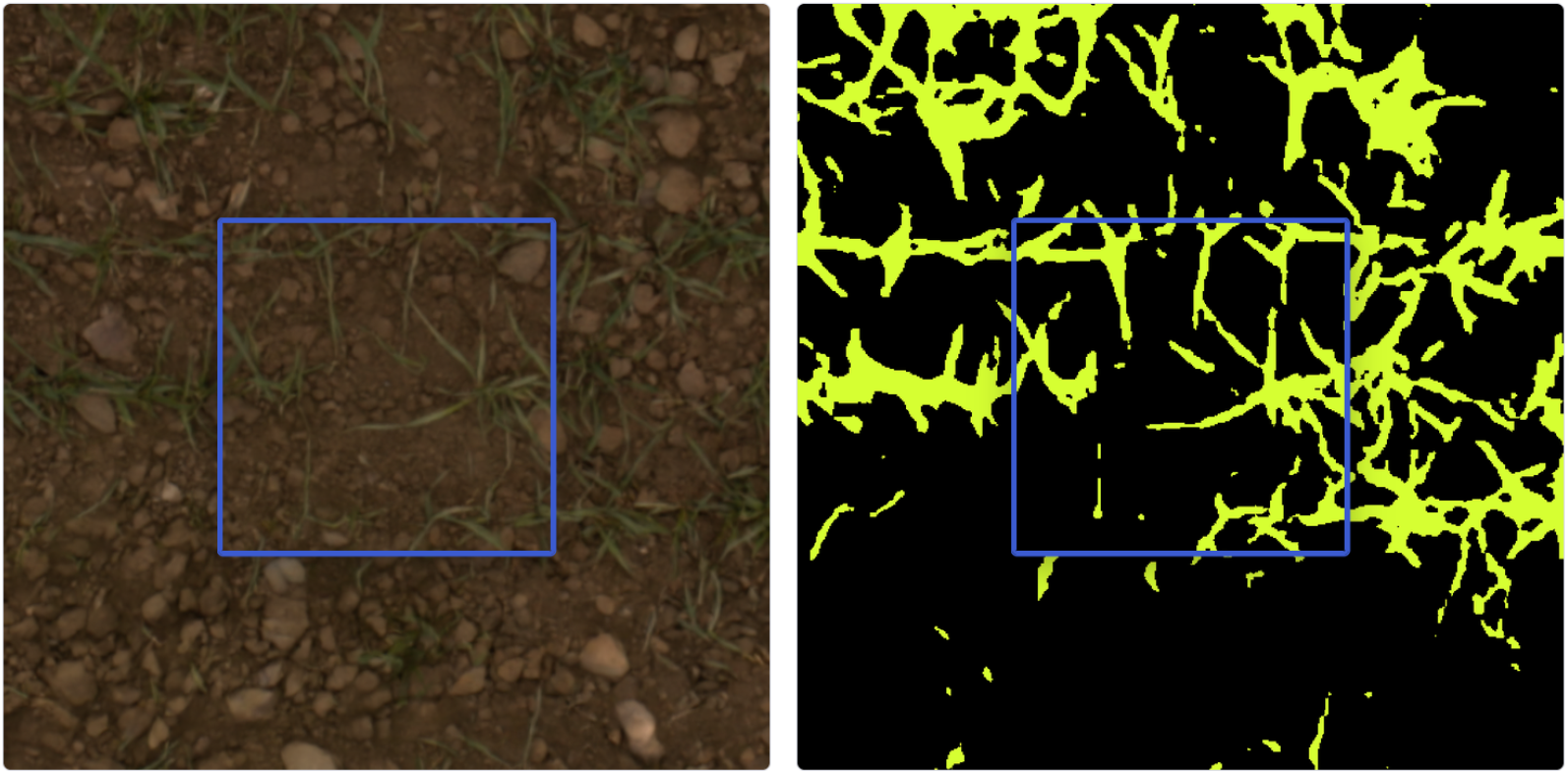
Progressive spatial outpainting from a single seed tile. Starting from one real image-mask tile (outlined in blue), OneGrow grows a larger canvas by iterated half-overlap outpainting, jointly generating the surrounding image (*left panel*) and organ mask (*right panel*). Each panel is one assembled canvas and is not split into reference and generated halves, unlike Fig. 3. Each outward window is conditioned on its overlap with the already-generated canvas, and the fully-convolutional decoder renders the assembled canvas in a single pass. Masks use the colors of Fig. 2.

### 5.4 Temporal roll-out

The temporal roll-out (Section 3.7) generates an image sequence from a mask sequence, re-feeding each generated image as context for later frames. We show this qualitatively on a FIP1 growth sequence, where real images seed the roll-out and each later frame is generated from its mask and the preceding frames (Fig. 7).

**Figure 7:**
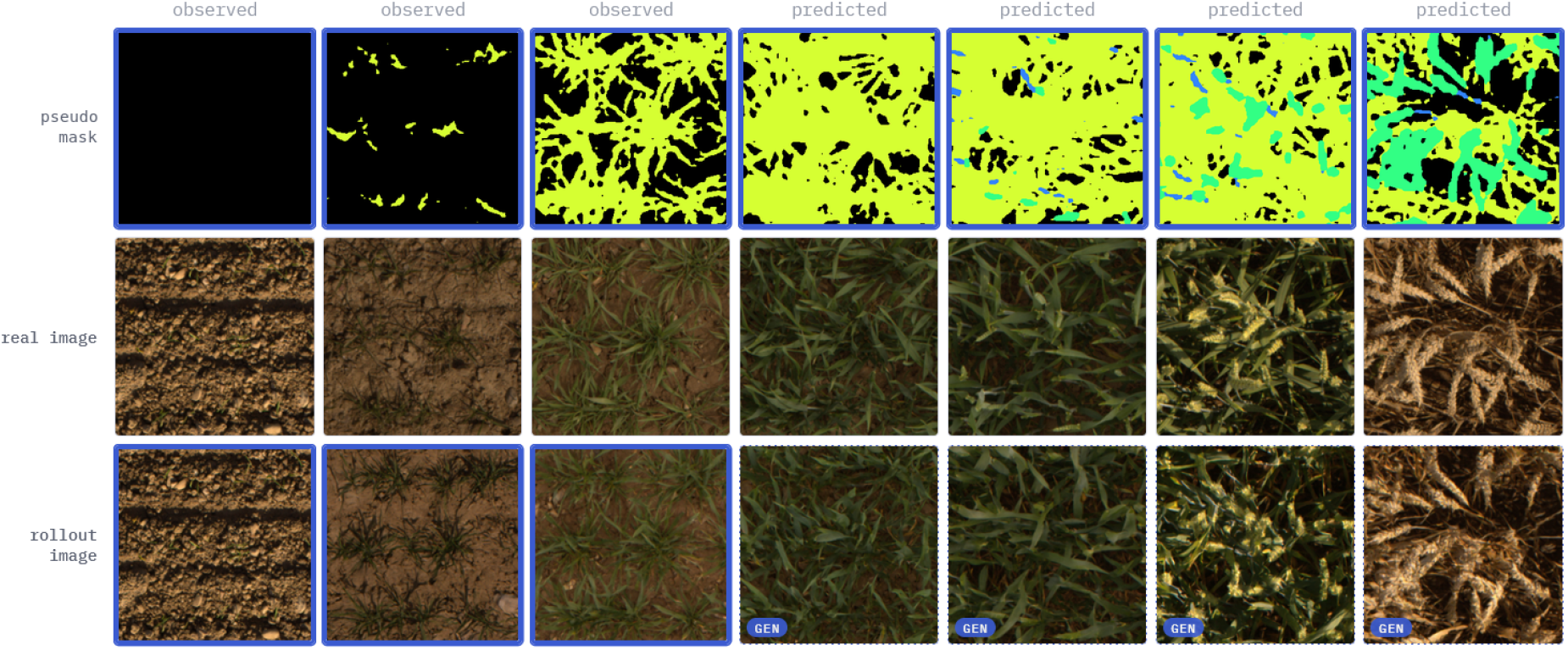
Qualitative temporal roll-out on a FIP1 growth sequence. Each column is one time step, ordered left to right. *Top:* the teacher pseudo-mask, revealed for every frame (blue border). *Middle:* the real image, shown for reference. *Bottom:* the roll-out. The first three frames (*observed*, blue border) are real images that seed the roll-out. Each later frame (*predicted*, marked gen) is generated from its mask and the three preceding frames, so generated images re-enter as context as the window slides. Masks are shown as background (black), leaf (yellow-green), stem (blue), and head (green).

## 6 Conclusion

We introduce OneGrow, a unified latent flow-matching model for wheat images and organ masks over time. A shared image-mask representation and per-patch reveal mechanism support tasks such as image-to-mask segmentation, mask-to-image synthesis, and forecasting. A sliding-window roll-out conditions each new image on its mask and preceding image context. Training combines a diverse single-frame source with a multi-year temporal source using pseudo-labels from a pretrained segmentation model. In its mask-to-image setting, OneGrow places organs where the conditioning mask specifies, reaching a cycle-consistency mean intersection-over-union of 0.749, on par with the 0.740 the GWFSS challenge winner reaches on real images.

### Limitations

The pseudo-labels inherit the biases of the pretrained segmentation model, particularly for thin stems. OneGrow encodes images and masks with a frozen, general-purpose autoencoder that we do not fine-tune, so it learns no latent representation of its own that could be analyzed for downstream comparisons. The model is trained only on wheat, so transfer to other crops and species remains untested. The mask-conditioned time-series generation that motivates the method is so far shown only qualitatively and needs quantitative validation. OneGrow does not match the challenge winner on image-to-mask accuracy, as expected. Image-to-mask segmentation is not a target task and is down-weighted in the training distribution, which emphasizes mask-conditioned synthesis and forecasting. OneGrow also learns from pseudo-labels produced by that same model, so its masks approximate the teacher rather than the human annotations used for evaluation. OneGrow is also more expensive at inference than a dedicated segmenter, since it generates through iterative sampling. It is trained under a limited compute budget, and larger-scale training may narrow the remaining quality gap.

### Applications and outlook

Because OneGrow pairs realistic imagery with known organ masks, the generated image-mask pairs could augment training data for segmentation and related recognition tasks [18, 55], covering growth stages and imaging conditions that are under-represented in real data. The temporal roll-out could also translate mask sequences from plant models [2] into realistic image sequences. Such models, and the temporal platform itself, also expose growth-influencing variables such as genotype and environment [15, 41]. These could enter OneGrow as further register-token conditioners alongside the acquisition domain and photometric statistics it already uses, giving explicit control over the synthesized development.

## Acknowledgements

The authors thank Achim Walter (ETH Zürich) for providing the research environment in which this work was carried out.

## A Additional Qualitative Results

This appendix shows reveal configurations not covered in Section 5.3. The figure conventions of Section 3.5 apply. Content outlined in blue is revealed as clean context, and every generated cell is split down the middle into the held-out reference (left half) and the generated content (right half).

### Joint spatial completion on FIP1

Figure 8 reveals only a central image patch of each frame, without any mask. OneGrow then generates the surrounding image and the full organ mask jointly, which distinguishes this task from the mask-guided completion of Fig. 5.

**Figure 8:**
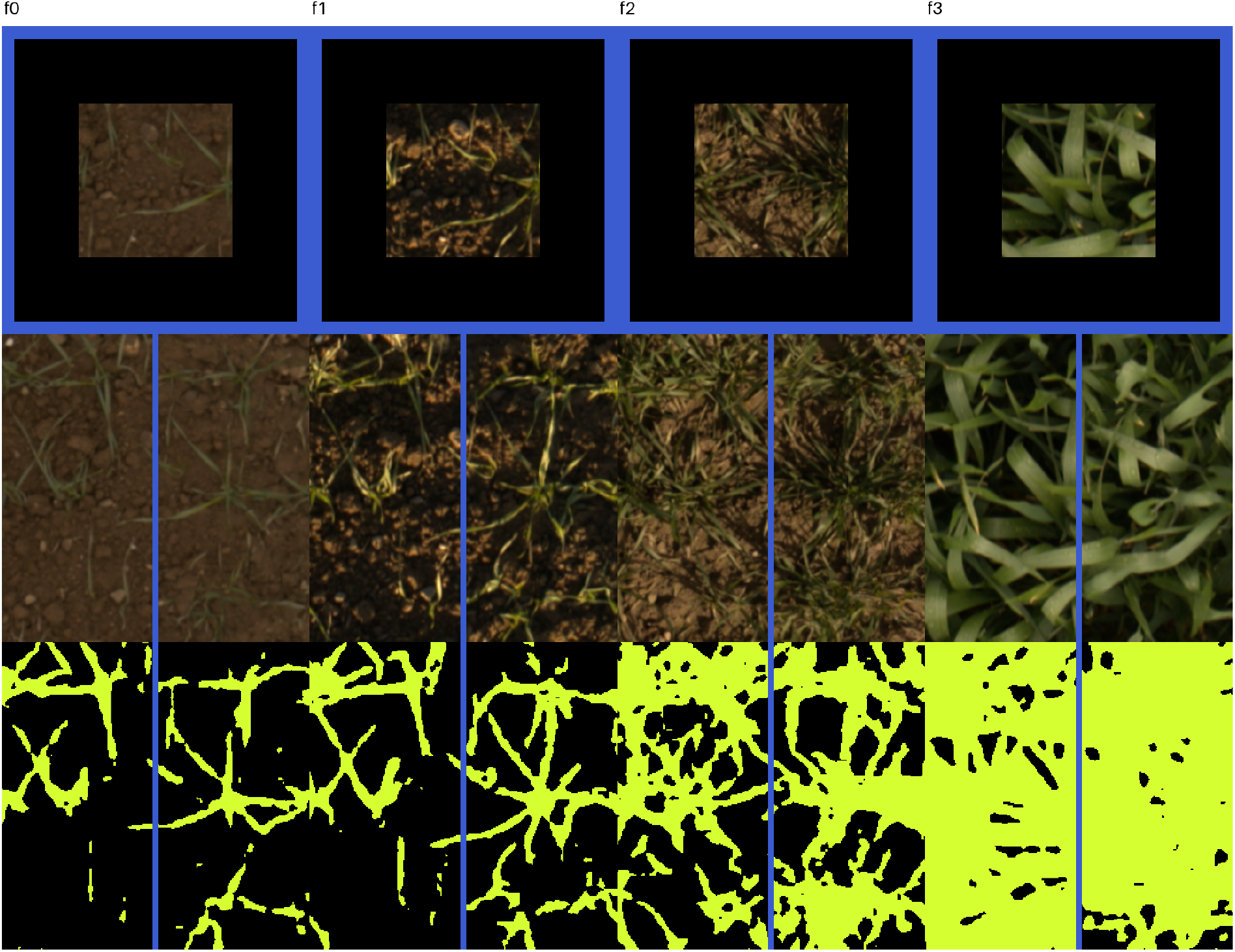
Joint spatial completion on a FIP1 growth window. Only a central image patch is revealed (*top row*), and OneGrow jointly generates the surrounding image (*middle row*) and the full organ mask (*bottom row*). Cells in the two generated rows are split down the middle into the held-out reference (left half) and the generated content (right half). No mask is given here, which distinguishes this task from Fig. 5.

#### Single-frame tasks on GWFSS

GWFSS contributes single-frame windows, so the same reveal configurations apply to one frame at a time (Figs. 9 and 10). These images are later-stage canopies and contain stem pixels (blue) alongside background (black) and leaf (yellow-green).

**Figure 9:**
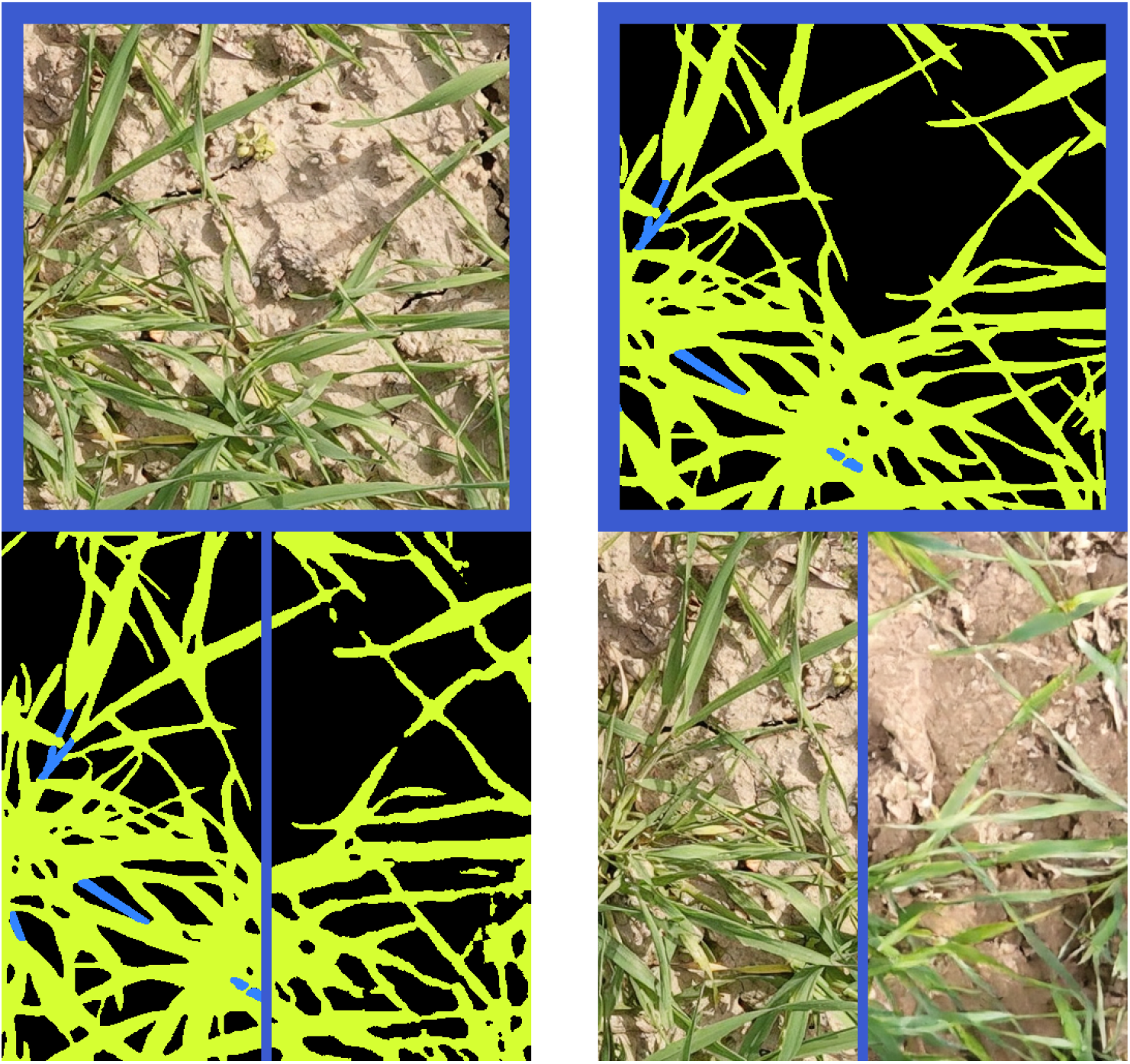
Single-frame tasks on GWFSS. *Left panel:* image-to-mask segmentation, where the image is revealed (top row) and the organ mask is generated (bottom row). *Right panel:* mask-to-image synthesis, where the mask is revealed (top row) and the image is generated (bottom row). Each generated cell is split down the middle into the held-out reference (left half) and the generated content (right half).

**Figure 10:**
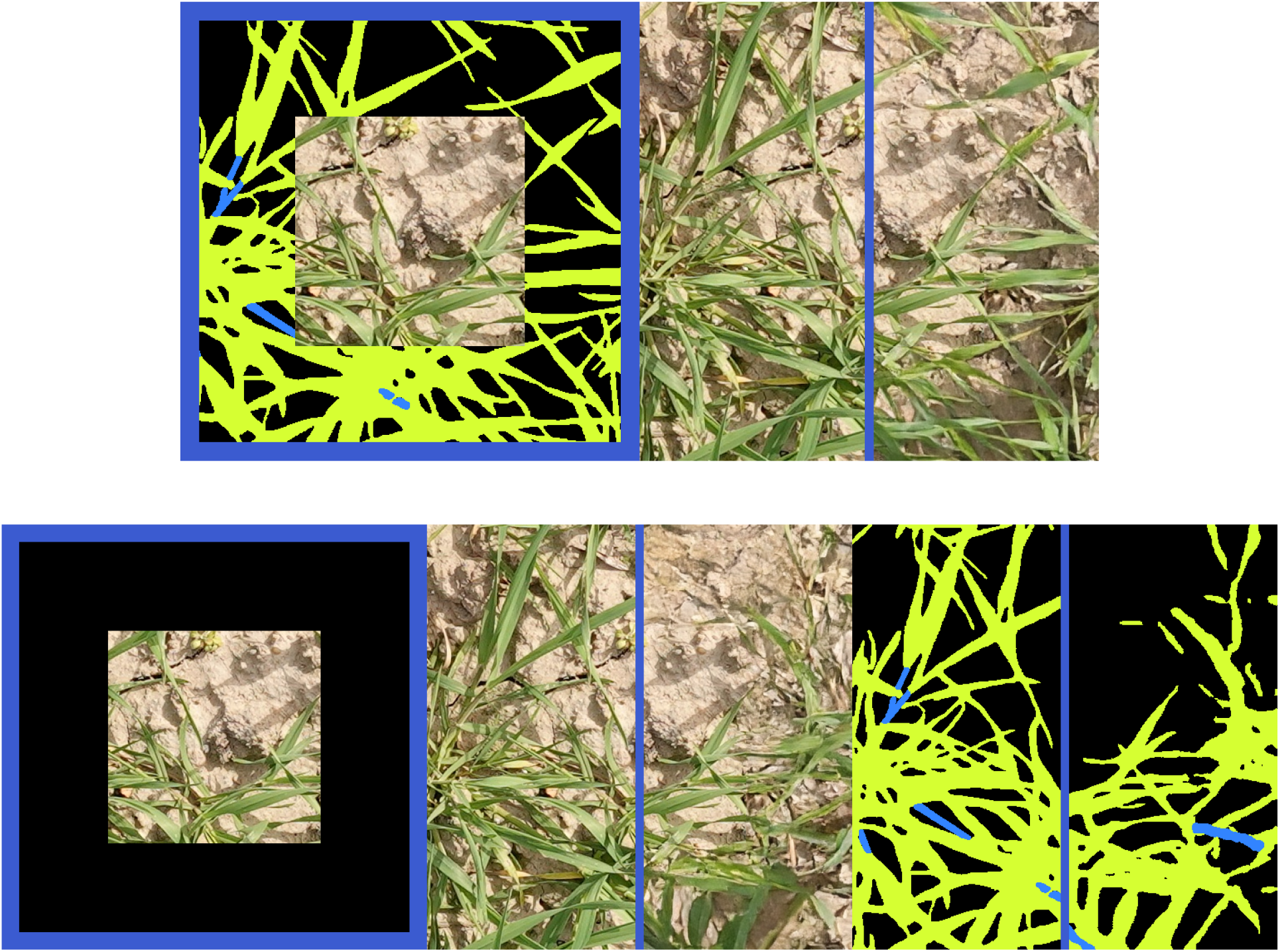
Spatial completion on GWFSS. Cells run left to right within each row. *Top:* mask-guided completion, where the full organ mask with a central image patch composited into its center is revealed (first cell) and the surrounding image is generated (second cell). *Bottom:* joint completion, where only the central image patch is revealed (first cell) and OneGrow generates both the surrounding image (second cell) and the organ mask (third cell). Each generated cell is split down the middle into the held-out reference (left half) and the generated content (right half).

